# Genetic and physiological insights into the diazotrophic activity of a non-cyanobacterial marine diazotroph

**DOI:** 10.1101/2022.07.13.499682

**Authors:** Aurélie Joublin-Delavat, Katia Touahri, Pauline Crétin, Amandine Morot, Sophie Rodrigues, Bruno Jesus, Florian Trigodet, François Delavat

## Abstract

Nitrogen (N_2_) fixation, or diazotrophy, supports a large part of primary production in oceans. Culture-independent approaches highlighted the presence in abundance of marine non-cyanobacterial diazotrophs (NCD) but their ecophysiology remains elusive, mostly because of the low number of isolated NCD and because of the lack of available genetic tools for these isolates. Here, a dual genetic and functional approach allowed unveiling the ecophysiology of a marine NCD affiliated to the species *Vibrio diazotrophicus*. Physiological characterization of the first marine NCD mutant obtained so far was performed using a soft-gellan assay, demonstrating that a Δ*nifH* mutant in not able to grow in nitrogen-deprived media. Furthermore, we demonstrated that *V. diazotrophicus* produces a thick biofilm under diazotrophic conditions, suggesting biofilm production as an adaptive response of this NCD to cope with the inhibition of nitrogen-fixation by molecular oxygen. Finally, the genomic signature of *V. diazotrophicus* is essentially absent from metagenomic data of *Tara Ocean* expeditions, despite having been isolated from various marine environments. We think that the genetically tractable *V. diazotrophicus* strain used in this study may serve as an ideal model to study the ecophysiology of these overlooked procaryotic group.

## Introduction

Nitrogen is a macro-element essential for life, as it is found in many biological molecules (*i.e*. nucleic acids, proteins). The atmosphere is composed by 78% of dinitrogen (N_2_) but this form cannot be assimilated by the vast majority of the organisms. The process of N_2_ fixation or diazotrophy is pivotal in oceanic ecosystems, since it makes the newly fixed nitrogen accessible to the entire plankton community, fueling primary production (Tang et al., 2019). Diazotrophy is now considered a key player in the biogeochemical cycles of nitrogen, but also of carbon, since it stimulates the biological pump, a mechanism absorbing a significant portion of the carbon dioxide emitted into the atmosphere and exporting it to the ocean seafloor. Diazotrophy is particularly crucial in tropical and subtropical area, where oceanic primary production is typically limited by the limited dissolved inorganic nitrogen (DIN) bioavailability (Capone et al., 2005). In addition, recent *in situ* quantification of N_2_ fixation demonstrated high diazotrophic activity in DIN-replete marine environments (Mulholland et al., 2019; Tang et al., 2019), showing the overall importance of diazotrophy in oceans.

Diazotrophy is an anabolic process restricted to some prokaryotic species (found both in *Bacteria* and *Archaea*). *Cyanobacteria* like the free-living *Trichodesmium* are known since decades to be important oceanic nitrogen fixers (Carpenter and Romans, 1991), accounting for at least 1.6 Tmol N in the tropical North Atlantic alone (Capone et al., 2005). Other *Cyanobacteria*, typically living in symbiotic association with photosynthetic microalgae also contribute significantly. These associations include diatom-diazotroph associations (DDA), where filamentous *Cyanobacteria* like *Richelia intracellularis* and *Calothrix rhizosoleniae* can be located either intra- or extracellularly to fix N2 and transfer the newly fixed nitrogen to sustain diatom growth (Nieves-Morion et al., 2020), receiving organic carbon from its host in return (Foster et al., 2022). DDA associations can account for 0.19 and 0.62 Tmol N per year in Atlantic and Pacific oceans, respectively (Foster et al., 2011). Another example includes the unicellular cyanobacterial lineage UCYN-A, which associate with haptophytes and fixes N_2_ even under nitrogen-replete conditions (Mills et al., 2020) and which significantly contribute to N_2_ fixation in oceans (Martinez-Perez et al., 2016).

Despite the tremendous data related to cyanobacterial N_2_-fixation in oceans, the global N budget in frequently considered unbalanced, as the N loss by denitrification and anammox largely exceeds N input (Landolfi et al., 2018). This suggests that alternative players are missing to equilibrate the N budget. More and more evidence show the presence in all oceans of previously overlooked nitrogen-fixers, not affiliated to the phylum *Cyanobacteria*. Indeed, culture-independent approaches based on the amplification or immunodetection of the *nifH* gene, a gene encoding a subunit of the nitrogenase, conserved among diazotrophs and considered as a biomarker for diazotrophy, have demonstrated the presence of marine Non-Cyanobacterial Diazotrophs (NCDs) (Zehr et al., 1998) in the water column attached to particles (undefined aggregates, faecal pellets) (Farnelid et al., 2019; Geisler et al., 2019) and in deep sea sediments (Kapili et al., 2020). Moreover, *Tara Oceans* expeditions provide metagenomic data from diverse oceanic locations, depths and size fractions, coupled with physical and chemical information (Karsenti et al., 2011; Planes et al., 2019). Analyses of the metagenomes allowed the reconstructions of MAGs (Metagenome-Assembled Genomes) from diverse diazotrophs, including many NCDs, found in abundance in diverse oceans (Delmont et al., 2018; Delmont et al., 2022). Metatranscriptomic analysis from *Tara Oceans* data showed that many NCDs express their *nifH* gene and are therefore active (Delmont et al., 2022).

From a biochemical point of view, diazotrophic activity faces a strong metabolic constraint under oxic conditions (Gallon, 1992). Indeed, O_2_ acts at different levels to block diazotrophy: it can (i) inhibit the expression of *nif* genes including *nifH* (Hubner et al., 1991; Dixon and Kahn, 2004), (ii) lead to irreversible damages of the nitrogenase (Gallon, 1992), and (iii) reversibly inhibit the nitrogenase activity (Goldberg et al., 1987). This O_2_-based inhibition of diazotrophy poses a metabolic conundrum, since many diazotrophs are also aerobes and therefore use O_2_ as a terminal electron acceptor for aerobic respiration. To solve this dilemma, terrestrial diazotrophs committed to symbiotic interactions with legume roots (and sometimes on stems (Bonaldi et al., 2011)) are engulfed within a plant organ called nodule. This nodule serves as a physical barrier, limiting O_2_ diffusion and creating micro-aerobic environments where both nitrogen fixation and respiration co-occur. Other terrestrial diazotrophs, like the free-living diazotroph *Azotobacter vinelandii* produces a cytochrome *bd* terminal oxidase, that consumes a high volume of O_2_, which leads to a rapid decrease in intracellular O_2_-tension, allowing nitrogen fixation in overall aerobic environments (Kelly et al., 1990; Poole and Hill, 1997). Additionally, *A. vinelandii* secretes an alginate capsule, which thickness increases with O_2_ tension and which serves as a barrier for O_2_ diffusion into the cells (Sabra et al., 2000).

In addition to the O_2_ tension found in oceans, marine *Cyanobacteria* face another challenge when regarding nitrogen fixation, since many of them are photosynthetic and release O_2_ as a byproduct of oxygenic photosynthesis. Marine *Cyanobacteria* have therefore also developed various strategies to solve this metabolic constraint. Free-living filamentous *Cyanobacteria* like *Anabaena* sp. (Zhang et al., 2006), but also those living in symbiotic associations with photosynthetic diatoms like *R. intracellularlis* and *C. rhizosoleniae* (Foster et al., 2011) phenotypically differentiate into distinct subpopulations: when DIN becomes limiting, some cells within the filament specialize in nitrogen-fixing factories. These cells, called heterocysts have a modified envelope containing additional layers (a glycolipid and a polysaccharide layer) which restricts O_2_ diffusion in the cell cytoplasm (for a review, see (Flores et al., 2019)). Heterocysts commit to a dedicated developmental program (Flores et al., 2019), also marked by the loss of ability to perform oxygenic photosynthesis, which is restricted to the neighboring vegetative cells (Zhang et al., 2006). Molecular exchanges of nitrogen and carbon compounds therefore occur between the vegetative cells and the heterocysts to sustain the growth of the filament (Nieves-Morion et al., 2021). As an alternative to this spatial regulation of diazotrophy, some unicellular *Cyanobacteria* like *Crocosphaera* perform oxygenic photosynthesis during the day and nitrogen fixation during the night (Tuit et al., 2004). Finally, non-heterocyst diazotrophs may protect their nitrogenase from O_2_ by the production of hopanoid lipids (Cornejo-Castillo and Zehr, 2019).

In contrast, while strategies deployed by marine cyanobacterial diazotrophs to perform nitrogen fixation in oxic environment are documented, knowledge on those for marine NCD remain understudied (Bombar et al., 2016). This knowledge gap is due to the limited number of NCD isolated from marine environments (Farnelid et al., 2014; Martinez-Perez et al., 2018). Additionally, their physiology remains elusive because, to our knowledge, no genetic mutant of marine NCD has been obtained so far, hampering an in-depth physiological characterization of marine NCD.

*Vibrio diazotrophicus* is a marine bacterium, firstly isolated from gastrointestinal tract of sea urchins (Guerinot and Patriquin, 1981). It can be found in marine and estuarine environments (Guerinot et al., 1982) as well as in sediments (Castillo et al., 2018) and in enriched marine phytoplankton exometabolites (Fu et al., 2020). The type strain *V*. *diazotrophicus* NS1 (ATCC 33466) was shown by acetylene reduction assay (ARA) to produce nitrogenase and fix N_2_ (Guerinot and Patriquin, 1981). The genome of *V. diazotrophicus* NS1 is available (accession number BBJY00000000.1, no associated publication or genomic characterization) and other isolates have been fully sequenced (Castillo et al., 2018). However, besides the description of its isolation in 1981 and its taxonomical characterization one year latter (Guerinot et al., 1982), no further physiological or genetic characterization of any member of this species has been performed. Because of its presence in various oceanic environments and because many genetic and molecular tools are available for this genus (Delavat et al., 2018; Morot et al., 2021), we think that *V. diazotrophicus* can constitute an ideal model to assess the ecophysiology of marine NCD and their diazotrophic activity.

In this manuscript, we report the physiological description of *V. diazotrophicus* NS1, together with the construction of the first genetic mutant of a marine NCD, impaired in nitrogen-fixation. Importantly, we showed that under nitrogen-deficient conditions, *Vibrio diazotrophicus* produces more biofilm, suggesting the role of biofilm as a barrier of O_2_ diffusion.

## Material and Methods

### Strains and culture conditions

The strains, plasmids and primers used in this study can be found in Table S1, S2 and S3, respectively. Unless otherwise stated, *V. diazotrophicus* NBRC 103148 and *Escherichia coli* were grown in LB at 30°C and 37°C, respectively. The nitrogen-deficient media used in this study is a modified diazotrophic medium for *Vibrio* (MDV), composed similarly to (Guerinot and Patriquin, 1981), with the following modifications: solution 1 was prepared with NaCl 0.3 M, MgSO_4_, 7H_2_O (0.05 M), CaCl_2_, 2H_2_O (0.01 M), KCl (0.01 M), Tris (0.05 M) adjusted to pH 7.5, before the addition of 0.3 g.l^−1^ yeast extract. Solution 2 was prepared in 500 ml H_2_O with 0.005 g Na_2_MoO_4_, 2H_2_O, 40 g glucose, 1 ml of a 3 g.l^−1^ FeNaEDTA,3H_2_O stock solution. Solution 3 was prepared in 166 ml H_2_O with 0.2 g KH_2_PO_4_ and 0.8 g K_2_HPO_4_ adjusted to pH 7.2. All 3 solutions were autoclaved, allowed to cool separately, and MDV was prepared by mixing 333 ml of solution 1, 500 ml of solution 2 and 166 ml of solution 3. 1 ml of a vitamin cocktail (stock solution containing 1 mg.l^−1^ biotin, 1 mg.l^−1^ vitamin B12 and 0.2 g.l^−1^ thiamin-HCl) was subsequently added to 1 l MDV.

If necessary, trimethoprim (Trim, 10 μg.ml^−1^), kanamycin (Km, 50 μg.ml^−1^), diaminopimelic acid (DAP, 0.3 mM), glucose (0.3 g.l^−1^), L-arabinose (L-ara, 0.2%) or agar (1.5%) were added to the media.

Doubling times were determined by spectrophotometry using the TECAN Infinite M1000. Briefly, one colony of each studied strain was inoculated in LB+Trim and incubated overnight. The next morning, cells were washed 3 times to remove old LB from the overnight culture, OD_600nm_ was adjusted to 0.01 with fresh LB, and 200 μl were used to inoculate 4 wells from a polystyrene plate (Dutscher, ref 330035). Plate was left in the TECAN instrument set at 30°C without shaking, and OD_600nm_ was measured every 30 minutes for 48 hours. Doubling times were calculated from the exponential growth phase of each culture.

### Strain constructions and DNA techniques

Standard procedures were used for all molecular techniques, following the reagent suppliers recommendations. Deletion of *nifH* was performed by double homologous recombination between a suicide plasmid and the chromosome of *V. diazotrophicus*, following a previously established protocol (Morot et al., 2021), with some adaptations: *V. diazotrophicus* was grown overnight at 30°C (it does not grow at 37°C), cointegrates were incubated in LB+L-ara liquid before plating on LB+L-ara, and the temperature used to incubate the filter-deposited LB+DAP plates was 30°C. The suicide plasmid pFD114 (Table S2) was constructed by amplifying around 750 bp of the upstream and downstream regions of the *nifH* gene of *V. diazotrophicus* (BBJY01_570151), using primers 210210 to 210213 (Table S3). Correct deletion of *nifH* was verified using primers 210501 and 210502.

Complementation of *nifH* was achieved by amplifying the *nifH* gene and its dedicated promoter, using primers 210204 and 210205 (Table S3). The P*_nifH_-nifH* amplicon was cloned within the pGEM-T vector (Promega), verified by sequencing and cloned within pFD086 (Morot et al., 2021) to replace P_*lac*_-gfp, using BamHI and XhoI. The resulting pFD120 plasmid was introduced by biparental mating as described above, using β3914 containing pFD120 as a donor strain. All *in silico* plasmid maps are available upon request.

Electroporation assays were performed following an already published method (Delavat et al., 2018), except the growing temperature of *V. diazotrophicus*, set at 30°C.

### Soft-gellan Assay

This medium was prepared as follows (for one tube): 6.7 ml of solution 1 without yeast extract plus 2 mg gellan gum were distributed in 18 cm glass tubes (Dutscher, ref 508232); 10 ml of solution 2 containing 3 mg gellan gum in a separate bottle; 3.33 ml of solution 3 containing 1 mg gellan gum in a separate bottle. All solutions were autoclaved separately, allowed to cool to 45°C, mixed in the glass tube, gently homogenized by inversion, before the addition of 20 μl of vitamin and Trim. If necessary, 20 μl of NO_3_^−^ (stock solution NaNO_3_ at 46.67 g.l^−1^) were added together with vitamins and Trim after autoclave. If necessary, 20 mg NH_4_Cl were added in the 3.33 ml of solution 3 before autoclave.

*V. diazotrophicus* containing the empty plasmid pFD085 and *V. diazotrophicus* Δ*nifH* containing either pFD085 or pFD120 were grown overnight in LB+Trim. 1 ml of overnight cultures was subsequently washed with 1 ml nitrogen-free MDV (without yeast extract), and pellets were resuspended with 500 μl nitrogen-free MDV. 100 μl of these washed suspensions were used to inoculate the soft-gellan tubes and tubes closed with a sterile blue rubber stopper, mixed by inversion and before being sealed with an aluminum capsule. Tubes were left for incubation at room temperature for 48 hours. The experiment was performed in triplicates.

Dissolved O_2_ concentrations were measured with a retractable fiber oxygen microsensor (tip diameter 50-70 μm, OXR50-OI, Pyro Science GmbH, Germany). The oxygen microsensor was connected to a FireSting-O_2_ fiber-optic oxygen meter (FireSting-O_2_, Pyro Science GmbH, Germany) and signals were recorded with a PC via the software Pyro Oxygen Logger (v3.319, Pyro Science GmbH, Germany). The sensor was linearly calibrated in distilled water by measuring in aerated water (100%) and in the same medium, in which 200 μl of Hcl-cysteine (5 %) was added after media preparation to reduce any trace of O_2_ (0%). The position of the sensor was controlled by a micromanipulator (MU1, Pyro Science, GmbH, Germany). Measurements were done before the medium-air interface and at the location of the bacterial ring.

### Biofilm production in microplates

Strains carrying either pFD085 or pFD120 were inoculated from a single colony in 5 ml LB+Trim and incubated overnight at 30°C with shaking. Part of the cultures were washed 3 times with LB, diluted to an initial OD_600nm_ of 0.01 with LB+Trim, and 200 μl of this OD-adjusted suspension were inoculated in quadruplicate in a 96-polystyrene-well microplate (Dutscher, ref 330035). The remaining overnight cultures were washed 3 times with MDV, adjusted to an initial OD_600nm_ of 0.01 with MDV+Trim, with MDV+Trim containing 46,67 mg.l^−1^ NO_3_^−^ or with MDV+Trim containing 20 mg.l^−1^ NH_4_Cl, and 200 μl of these OD-adjusted suspensions were inoculated in quadruplicate in the same microplate. Plates were placed in a microplate reader (TECAN Infinite M1000) for 48 hours at 30°C without shaking, and OD_600nm_ was measured every 30 minutes. After 48 hours, plates were treated following an already established protocol (Morot et al., 2021). Biofilm production was corrected by dividing the OD_550nm_ measured after crystal violet staining by the maximal OD_600nm_ measured during the growing time of the corresponding well.

### Biofilm production on flowcells

*V. diazotrophicus* biofilms were grown at 30°C under hydrodynamic conditions in a three-channel flow cell (1 × 4 × 44 mm; Biocentrum, DTU, Denmark (Pamp et al., 2009)). The flow system was assembled, prepared and sterilized as described by Tolker-Nielsen and Sternberg (Tolker-Nielsen and Sternberg, 2011). The substratum consisted of a microscope glass coverslip (24 × 50 mm; KnittelGlasser, Braunschweig, Germany). Each channel was inoculated with 250 μl of an overnight LB+Trim culture of *V. diazotrophicus* pFD086 diluted (after 3 washing steps in MDV) to an OD_600nm_ of 0.1 in MDV. A 2h-attachment step was performed without any flow of medium. Then a 5 ml.h^−1^ flow of MDV was applied for 24 h using a Watson Marlow 205U peristaltic pump (Watson Marlow, Falmouth, UK). Biofilms were observed by monitoring the GFP fluorescence with a LSM 710 Confocal Laser Scanning Microscope (Zeiss, Oberkochen, Germany) using a 40x oil immersion objective. GFP was excited at 488 nm and fluorescence emission was detected between 500 and 550 nm. Images were acquired at intervals of 1 μm throughout the whole depth of the biofilm. ZEN 2.1 software (Zeiss, Oberkochen, Germany) was used for visualization and image processing. Quantitative analyses of image stacks were performed using COMSTAT software (http://www.imageanalysis.dk/) (Heydorn et al., 2000). Analysis was performed from 3 biological replicates (independent experiments) and 8 technical replicates, every replicate comprising 3 to 5 stacked images.

### Statistical analyses

All statistical analyses were performed using GraphPad Prism; *p* values <0.05 were considered statistically significant.

#### Phylogenomics, pangenomics and metagenomics detection of V. diazotrophicus NS1

We used anvi’o 7.1 (Eren et al., 2021) and the contigs snakemake (Koster and Rahmann, 2018) workflow with the heterotrophic bacterial diazotroph MAGs from Delmont et al. (Delmont et al., 2022). Briefly, the workflow created a contigs database with “anvi-gen-contigs-database,” which used Prodigal version 2.6.3 (Hyatt et al., 2010) to identify open reading frames. It used “anvi-run-hmm” to detect the single-copy core genes from bacteria (n = 71, modified from (Lee, 2019)), ribosomal RNAs (rRNAs) (n = 12, modified from https://github.com/tseemann/barrnap). For the phylogenomic tree of *V. diazotrophicus* NS1 and the MAGs from (Delmont et al., 2022), we selected 16 bacterial SCGs present in at least 40 out of the 41 genomes, which we concatenated and aligned using Muscle v3.8.1551 (Edgar, 2004) and used FastTree 2.1.11 (Price et al., 2010) to generate a tree. We used “anvi-interactive” to visualize the phylogenomic tree.

To compare *V. diazotrophicus* NS1 with publicly available genomes of the same species, we used “ncbi-genome-download” (https://github.com/kblin/ncbi-genome-download) to get all *V. diazotrophicus* genomes from GenBank. We discarded two genomes for their very small size. We used anvi’o Pangenomics snakemake workflow to compare the gene content between the genomes. The workflow used Prodigal to identify open reading frames, DIAMOND v2.0.14 (Buchfink et al., 2015) to quantify the similarity between each pair of genes, the Markov Clustering Algorithm (MCL) to identify gene clusters, and Muscle v3.8.1551 to align amino acid sequences. We used “anvi-compute-genome-similarity” to compute the Average Nucleotide Identity (ANI) using pyANI v0.2.11 and visualized the pangenome using “anvi-display-pan”. We used the “functional homogeneity” index to filter for more divergent gene clusters (max functional homogeneity of 0.95), which we used to generate a *V. diazotrophicus* specific phylogenomic tree (n=99 genes) using the same approach as described above (Muscle v3.8.1551 and FastTree 2.1.11).

We used COG20 (Galperin et al., 2021), KOfam (Aramaki et al., 2020) and the InterPro website (Jones et al., 2014) to identify the nitrogen fixation associated genes. We extracted three loci from each genome using “anvi-export-loci” and the anchor genes nifH (K02588), nifU (K02594) and nifQ (K15790) and used the R package “gggenes” (https://wilkox.org/gggenes/) to visualize the gene synteny.

We have also used metagenomes from the TARA ocean project to recruit reads on to *V. diazotrophicus* NS1. Briefly, we use the “metagenomics” snakemake workflow in anvi’o and mapped metagenomics short-reads using Bowtie2 (Langmead and Salzberg, 2012) and summarized the results with `anvi-summarize`. We used the “detection” value to estimate the presence/absence of V. diazotrophicus NS1 in both the prokaryotic and large size fraction metagenomes.

## Results

### Optimization of genetic tools for V. diazotrophicus NS1

*V. diazotrophicus* NS1 is a marine strain, for which no genetic mutant has been described. Because this strain might serve as an ideal model for genetic and ecophysiological characterization of marine NCD, we sought to develop and optimize a genetic protocol for *V. diazotrophicus* NS1. To achieve this aim, we first applied and modified an electroporation protocol, originally developed for *Vibrio harveyi* and *Pseudoalteromonas* sp. (Delavat et al., 2018). We tested different variants of the protocol, by using either 1- or 2 mm electroporation cuvettes, modifying the voltage from 1500 to 2500 V in the presence or absence of 15% glycerol in the sucrose buffer (Delavat et al., 2018), from stationary-phase cultures grown in LB or LBS at 30°C. The best electroporation protocol yielded 81 CFU/μg DNA, and was obtained when the following protocol was applied: glycerol-containing buffer was used to wash LB-grown cultures, and the electroporation voltage was set at 2000 V in a 2 mm electroporation cuvette (data not shown). This demonstrates that, although being at low efficiency, electroporation can be applied for this strain.

In order to improve the efficiency of exogenous DNA transfer to *V. diazotrophicus* NS1, we also performed bi- and triparental mating as recently developed (Morot et al., 2021). Both conjugations were successful for transferring the replicative plasmid pFD086 to *V. diazotrophicus* NS1, giving rise to bright GFP-fluorescent cells (Figure 1). Finally, applying this protocol, we successfully inserted a mini-Tn7 in *V. diazotrophicus* NS1 containing the unstable pFD052 plasmid expressing the Tn7-transposase (Delavat et al., 2018), the latter being subsequently cured (data not shown).

**Figure 1.**
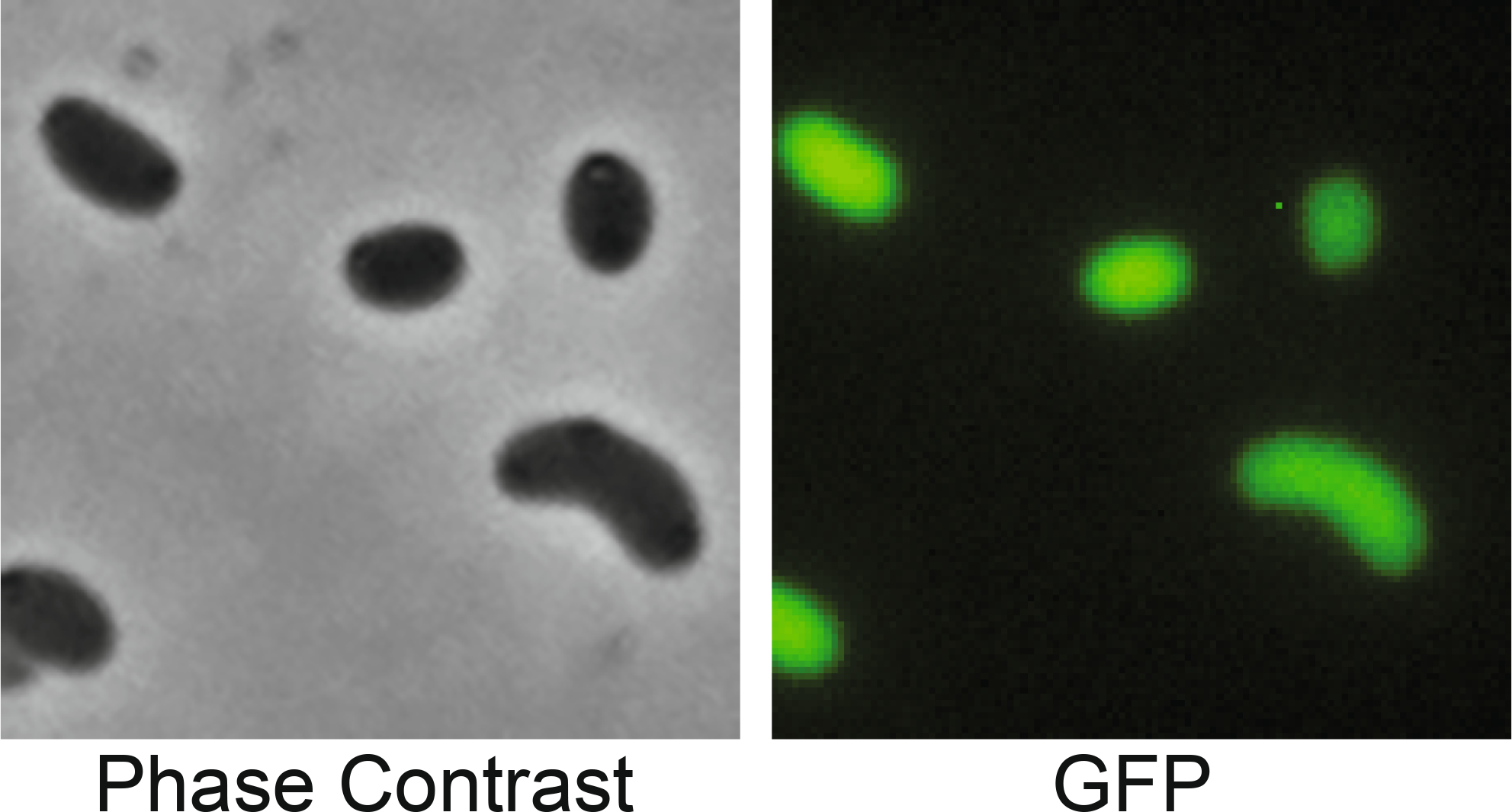
Microscopy picture showing the obtention of *V. diazotrophicus* carrying the GFP-expressing plasmid pFD086. Images were acquired from LB-grown overnight cultures.

### Construction of a nifH deletion mutant and physiological characterization

Unveiling the genetics and ecophysiology underlying diazotrophy in *V. diazotrophicus* requires the deletion of genes and the subsequent physiological characterization of the obtained mutants. In this study we focused on the deletion of the *nifH* gene (BBJY01_570151), applying an adapted protocol successfully used for *V. harveyi* ORM4 (Morot et al., 2021). Using this protocol, a clean *nifH* deletion mutant was obtained (Figure 2A). This mutant does not show any growth phenotype when growing in LB-rich medium (Figure 2B).

**Figure 2.**
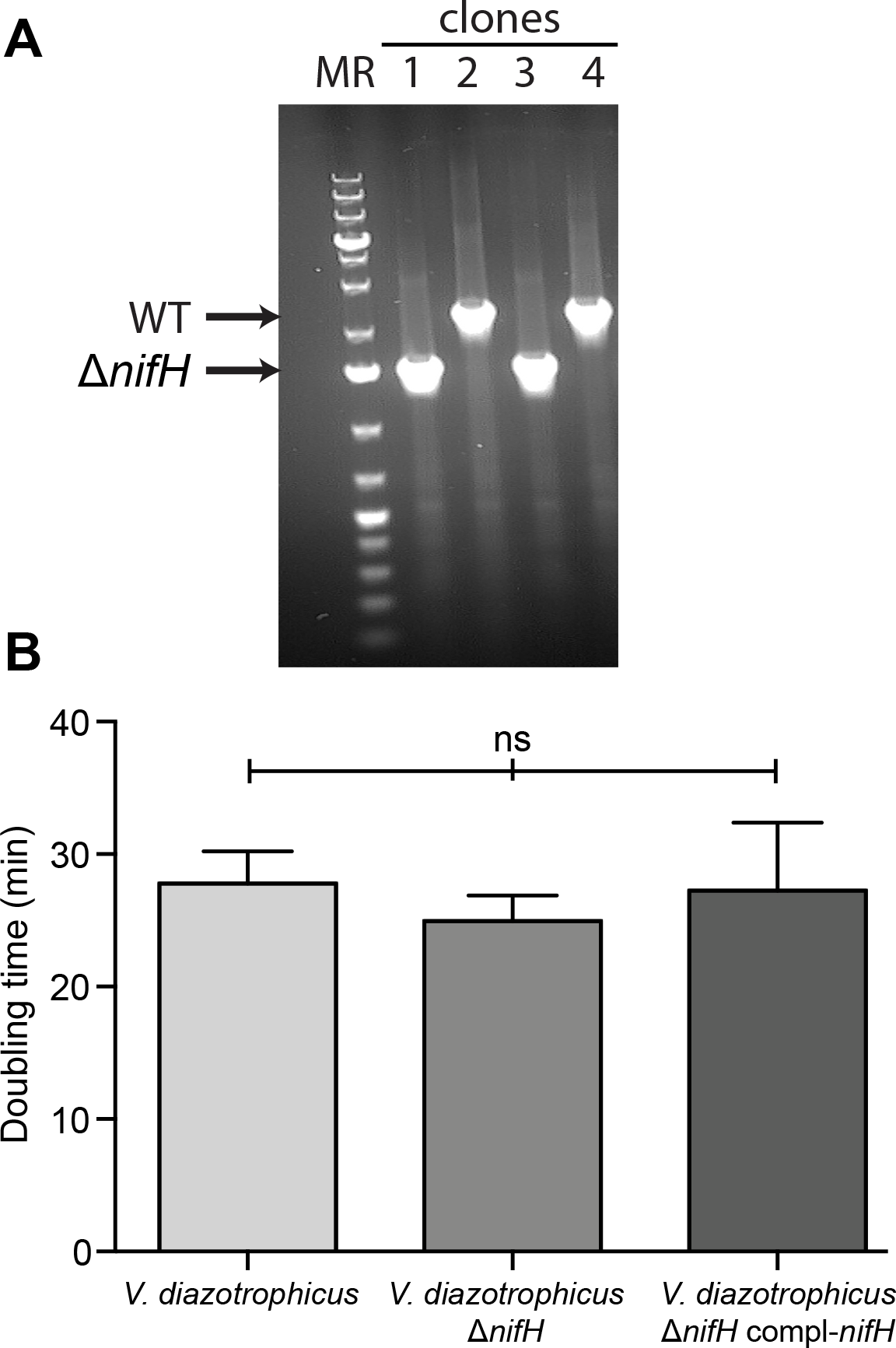
Genetic and physiological characterization of the *V. diazotrophicus* NS1 and the Δ*nifH*mutant. (A) Gel electrophoresis showing the deletion of *nifH* in *V. diazotrophicus* NS1 (lanes 2 and 4). (B) Generation time of *V. diazotrophicus* NS1 and derivative mutants. *V. diazotrophicus* NS1 and *V. diazotrophicus* NS1 Δ*nifH* contained the pFD085 plasmid and *V. diazotrophicus* NS1 compl-*nifH* corresponds to *V. diazotrophicus* NS1 Δ*nifH* containing pFD120. A one-way ANOVA with Tukey test was done to compare the different mutants. Depicted here are the mean+SEM of 4 replicates.

In order to characterize the diazotrophic activity of the *nifH* mutant, we developed a soft-gellan assay, characterized by the use of a nitrogen-deprived medium containing 0.3% gellan gum. Inoculating tubes sealed with a blue rubber stopper and an aluminum capsule (Figure 3) with *V. diazotrophicus* NS1 containing the empty plasmid pFD085 led to a growth of this strain throughout the entire tube height (with a higher biomass closer to the surface) when NH_4_^+^ is added. Importantly, in the absence of NH_4_^+^, a single ring is visible after 48h, located around 15 mm below the surface of the medium. Applying this assay for the *nifH* mutant containing pFD085 gave a similar pattern as the one of the wild type when grown with NH4+. However, in the complete absence of nitrogen, no band nor growth was detectable in the *nifH* mutant. Additionally, no band was visible when tubes were left for 2 more weeks, indicating that the observed phenotype does not correspond to a longer lag phase in this medium. *In trans* complementation of the *nifH* deletion mutant by the introduction by conjugation of a replicative plasmid containing the entire *nifH* gene and its P_*nifH*_ promoter restored the wild type phenotype, with the formation of a clear band located roughly 15 mm below the surface of the medium (Figure 3). We subsequently repeated the experiment, using NO_3_^−^ as an alternative bioavailable nitrogen source, and the results were identical (see Figure S1). Thus, we demonstrated that growth of *V. diazotrophicus* NS1 under nitrogen-deprived medium is dependent on an intact nitrogenase.

**Figure 3.**
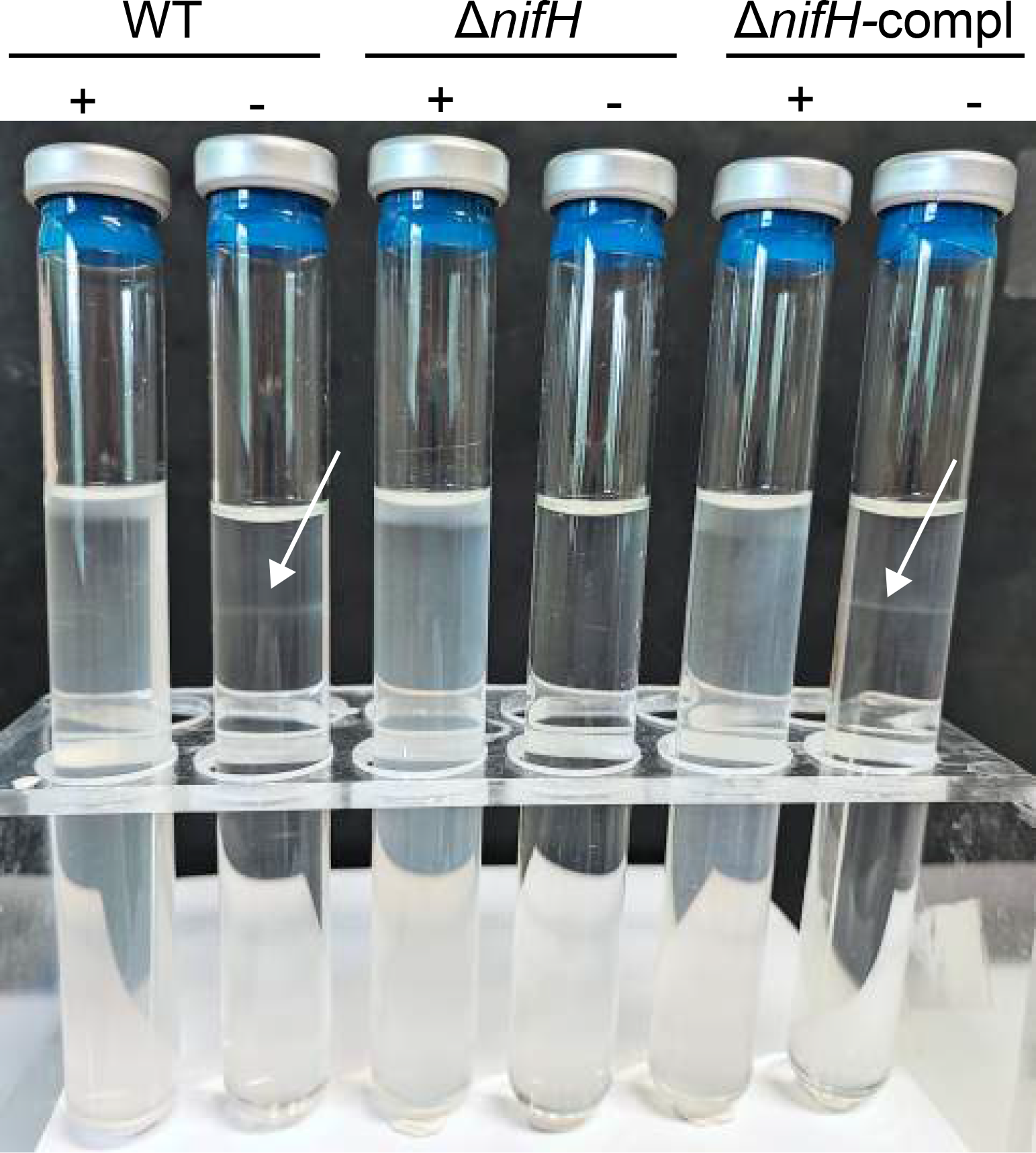
Soft-gellan assay demonstrating bacterial growth under appropriate microoxic conditions. The “+” and “−” indicate the presence or absence of NH_4_^+^, respectively. White arrows indicate the location of the growing ring. Note the absence of ring in the Δ*nifH* mutant and in the presence of NH_4_^+^. Similar results were obtained using NO_3_^−^ as a bioavailable inorganic nitrogen source (see Figure S1).

The O_2_ concentration in the soft-gellan tube was subsequently measured in the headspace and at the location (or corresponding depth) of the growing ring. Uninoculated soft-gellan tubes showed a high O_2_ concentration at both locations, with values ranging from 173.6 μM (sd 20.1) at a depth corresponding to the growing ring to 245.5 μM (sd 1.9) in the head space. In contrast, after 72 hours growth of *V. diazotrophicus* in nitrogen-free MDV, the O_2_ decreased to a concentration of 1.6 μM (sd 1.1) at the location of the ring, while this concentration was 238.8 μM (sd 1.4) in the headspace. To better approximate the O_2_ concentration at a depth where N_2_ fixation can occur, we quantified this concentration in 12 hours-inoculated tubes, where the growing ring is not yet formed. Under this condition, a O_2_ concentration of 14.8 μM (sd 7.7) is quantified at a similar depth as the one where the ring will form 2 days later. Thus, we demonstrated here that diazotrophy occurs under microaerobic conditions. Moreover, a subsequent soft-gellan experiment using *V. diazotrophicus* NS1 constitutively expressing GFP (thanks to the pFD086 plasmid it contains) showed that cells growing under microaerobic conditions within the ring are free-living and isolated, and don’t form aggregates (Figure S2).

### *V. diazotrophicus* produces more biofilm in MDV

In order to determine the possible regulatory mechanism deployed by *V. diazotrophicus* NS1 to avoid the nitrogenase inhibition by O_2_, we sought to quantify the ability of this strain to produce biofilm. Microplates containing either LB, MDV or MDV supplemented with either NO_3_^−^ or NH_4_^+^, plus trimethoprim were inoculated with either *V. diazotrophicus* NS1 (pFD085), *V. diazotrophicus* NS1 Δ*nifH* (pFD085) or *V. diazotrophicus* NS1 Δ*nifH* (pFD120), and left without shaking at 30°C. Neither *V. diazotrophicus* NS1 nor their derivatives do produce any biofilm when grown in LB. However, when grown in MDV, biofilm production significantly increased in the three tested strains (Figure 4A), and this biofilm was located at the bottom of the wells. Of note, the original MDV used for this experiment contains 100 mg.l^−1^ yeast extract, which provides low-but-necessary bioavailable nitrogen for the *nifH* mutant to grow, allowing direct comparison of the 3 tested strains. Importantly, when MDV was supplemented with NO_3_^−^ or NH_4_^+^, the biofilm production significantly decreased as compared to the one obtained in MDV medium. Additionally, we observed only subtle differences between the different tested strains, all three of them producing more biofilm in MDV as compared to LB and to MDV supplemented with either NO_3_^−^ or NH_4_^+^. This increase in biofilm formation in MDV suggests that in a nitrogen-limited medium, the production of biofilm by *V. diazotrophicus* NS1 might protect its nitrogenase from an excess in O_2_.

**Figure 4.**
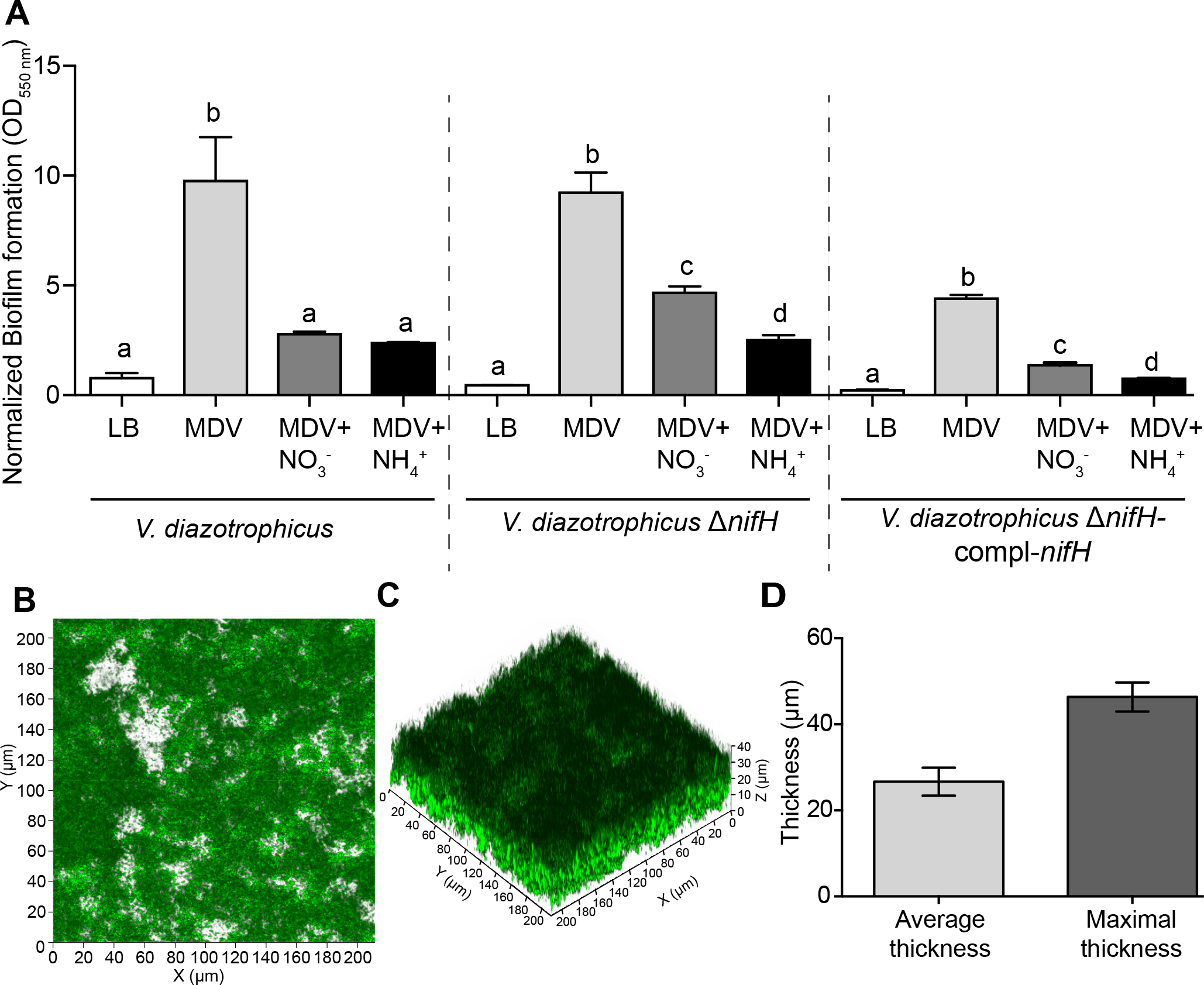
Biofilm formation characteristics of *V. diazotrophicus* NS1. (A) Corrected biofilm formation in *V. diazotrophicus* NS1 and derivative mutants in different media. *V. diazotrophicus* NS1 and *V. diazotrophicus* NS1 Δ*nifH* contained the pFD085 plasmid and *V. diazotrophicus* NS1 compl-*nifH* corresponds to *V. diazotrophicus* NS1 Δ*nifH* containing pFD120. Presented here are the mean+SEM of at least three replicates. Significance was assessed by one-way ANOVA with Tukey test for each mutant in the different media. (B) View of the top of the biofilm produced by *V. diazotrophicus* NS1 pFD086 after 24 hours of incubation in a flow cell. (C) Stacked microsopy image showing the thickness of the biofilm produced after 24 hours. (D) Average and maximal thickness of the biofilm produced by *V. diazotrophicus* NS1 pFD086 after 24 hours. See Figure S3 for images corresponding to timepoint 48 hours.

Furthermore, the thickness and biovolume of the biofilm produced in MDV by *V. diazotrophicus* were quantified by confocal microscopy, using flow cells. Under this condition, *V. diazotrophicus* produced an important biofilm (average biovolume of 11.7μm^3^.μm^−2^, Standard Error of the Mean SEM 1.64 μm). The thickness of the biofilm was also measured, reaching on average 26.65 μm (SEM 3.25 μm) and a maximal thickness of 46.35μm (SEM 3.34 μm) after 24 hours (Figure 4BCD). After 48 hours, these values increased even more, but the biofilm matrix was too dense and too heterogeneous, probably because of the maturation of the biofilm, causing early dispersal (Supplementary Figure S3).

### V. diazotrophicus is different from recently discovered heterotrophic bacterial diazotrophs

The genome of *V. diazotrophicus* NS1 was compared with the 40 Metagenome-Assembled Genomes (MAGs) of HBDs recently retrieved from *Tara ocean* metagenomes (Delmont et al., 2022). Expectedly, *V. diazotrophicus* NS1 falls in a group containing all MAGs affiliated to *Gammaproteobacteria* (Figure 5A). However, the closest MAG (called HBD_Gamma_02 here and in (Delmont et al., 2022)) seems to be affiliated to the order *Pseudomonales* and not to *Vibrionales*, and ANI comparison (value of 0.68, see Table S4) confirmed their phylogenomic distance. In addition, the genome of V. diazotrophicus NS1 was not detected in the Tara ocean metagenomes (max detection was 8% of the genome, Table S5). We subsequently compared the genomic relatedness of *V. diazotrophicus* NS1 to the genomes of 6 strains isolated from deep subsurface sediments of the Baltic Sea (Castillo et al., 2018), to the one isolated from phytoplankton exometabolite-enrichments (Fu et al., 2020) and to strain *V. diazotrophicus* 99A isolated in Pennsylvania (unpublished data). Pangenomic and phylogenomic analyses showed that *V. diazotrophicus* NS1 is closely related to 6 of these 8 strains (Figure 5B), with *V. diazotrophicus* 60.6B and HF9B standing a bit apart, possibly belonging to a new species (ANIb comparison below 0.95, see Figure 5B and Table S4). Of note, all of these nine *V. diazotrophicus* isolates carry a single copy of the *nifH* gene (Figure 5B and 5C) and a similar *nif* cluster (Figure 5C).

**Figure 5.**
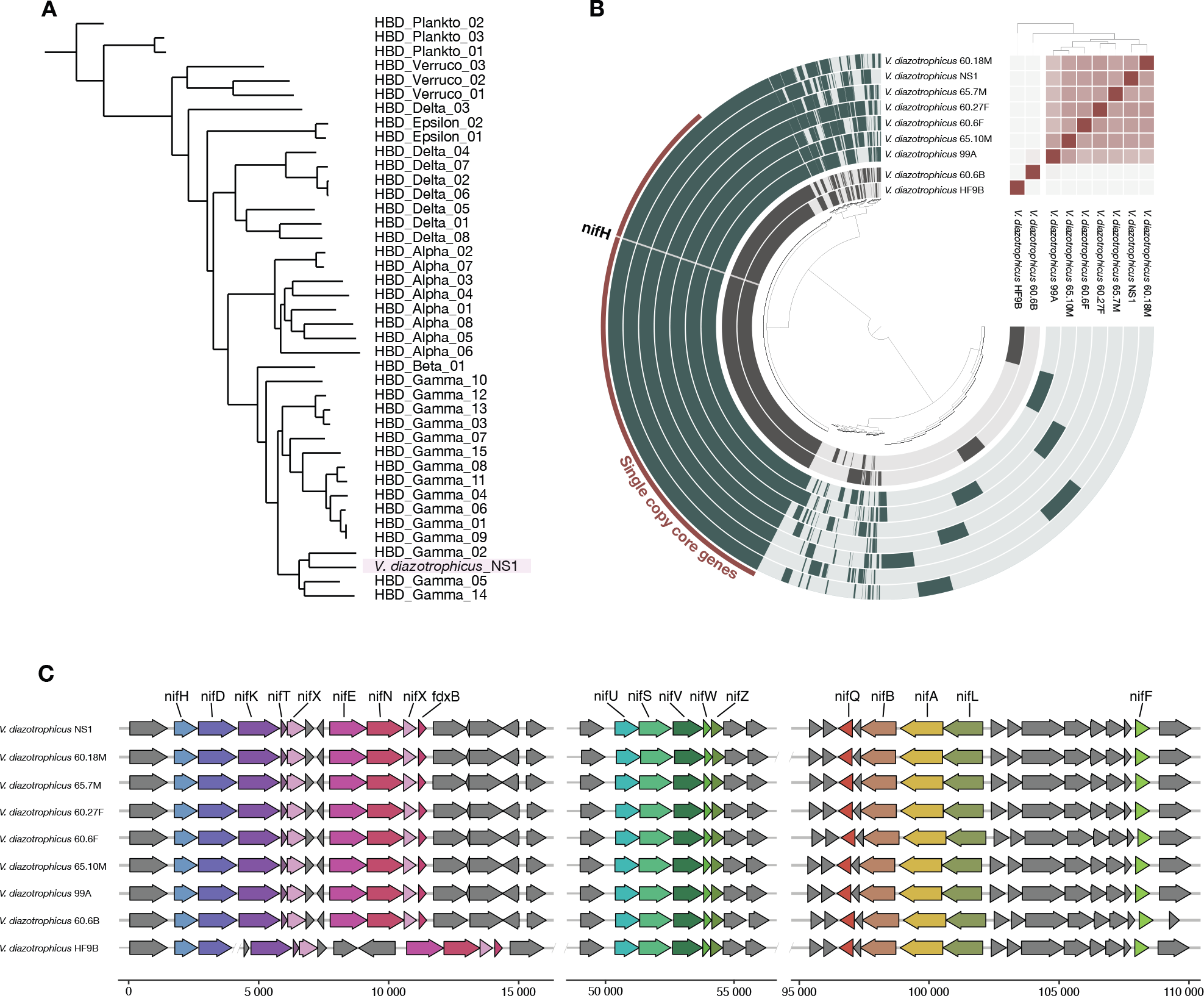
Comparative genomics of *V. diazotrophicus* NS1. (A) Phylogenomics tree of *V. diazotrophicus* NS1 and the 40 diazotroph MAGs from (Delmont *et al.*, 2022) using 16 single copy core genes occurring in at least 40 of the 41 genomes. We used FastTree to compute a phylogenomic tree and anvi’o (Eren *et al.*, 2021) for the visualization. (B) Pangenomics analysis of publicly available *V. diazotrophicus*. A total of 6,928 gene clusters are represented in the figure with each darker color in the genome’s layer indicating the presence of gene cluster in the corresponding genome. The red selection highlights the 2,884 single-copy core genes. The heatmap corresponds to the ANI computed between all genomes and it is scaled from below 0.95 ANI (white) to 1 ANI (dark red). We generated a phylogenomics tree (shown above the ANI heatmap) using a subset of set of 99 single copy core gene with a maximum functional homogeneity of 0.95 using anvi’o. (C) Conserved synteny of the *nif* gene locus across multiple *V. diazotrophicus* genomes. The scale is based on the total size of the locus in *V. diazotrophicus* NS1. The double bars indicates that the locus continues on a different contig. Acc. Num: 65.7M (POSL00000000), 65.10M (POSM00000000), 60.27F (POSK00000000), 60.18M (POSJ00000000), 60.6F (POSI00000000), 60.6B (POSH00000000), HF9B (JAATOR000000000) and 99A (PRJNA456207).

## Discussion

Nitrogen fixation is a major metabolism in many ecosystems, supplying living micro- and macro-organisms with otherwise scarce bioavailable nitrogen. In oceans, diazotrophy can account for half of the annual new production in euphotic zones in oligotrophic oceans (Karl et al., 1997). The diversity of marine diazotrophs has long been thought to be restricted to members of the phylum *Cyanobacteria*. However, culture-independent approaches, like those targeting the *nifH* gene or metagenomic and metatranscriptomic analyses from various oceanic samples demonstrated the presence in abundance of non-cyanobacterial diazotrophs (NCD, (Zehr et al., 1998; Delmont et al., 2018; Farnelid et al., 2019; Geisler et al., 2019; Kapili et al., 2020; Delmont et al., 2022)). Despite these observations, few marine NCD have been brought to culture (Farnelid et al., 2014; Martinez-Perez et al., 2018), hampering in depth understanding of their ecophysiology and contribution to global N budget in oceans. The goal of this study was to unveil the physiology of *V. diazotrophicus* NS1, a marine NCD isolated already 40 years ago (Guerinot and Patriquin, 1981) but surprisingly overlooked for in-depth analysis of its physiology.

The genome of *V. diazotrophicus* NS1 is available, and its analysis shows that it contains one *nif* cluster, separated in 3 chromosomal locations and spanning some 110 Kbp (Figure 5C). This cluster is found in all *V. diazotrophicus* strains whose genomes have been sequenced, with only minor differences. The presence of only one *nif* cluster is unlike other soil NCD like those belonging to the *Bradyrhizobium* genus, which can be endowed with 2 *nif* clusters (de Matos et al., 2021). In line with this observation, we showed here that the *nifH* gene of all sequenced *V. diazotrophicus* strains is part of the single-copy core gene fraction of the genome (Figure 5C), unlike many other *Bradyrhizobium* strains (de Matos et al., 2021).

Because *V. diazotrophicus* NS1 belongs to one of the few marine NCD isolated, we sought to develop and optimize genetic and molecular tools for this strain. Results show that *V. diazotrophicus* is genetically tractable, both by electroporation and conjugation. Electroporation efficiency was albeit low, as compared to the one reported for other *Vibrio* or *Pseudoalteromonas* species (Delavat et al., 2018). Improving the electroporation efficiency will require trials and errors, but having an already functional method reinforce its high potential for rapid introduction of exogenous DNA, strengthened by its functionality in other marine bacteria like *Algoriphagus machipongonensis* (Xu et al., 2022) or human pathogens like *Vibrio parahaemolyticus* (Soree et al., 2022). Conjugation was more efficient, both for the transfer of replicative plasmids and for the insertion of mini-transposons. Mating is however more time-consuming and requires efficient counter-selection of the donor. This counterselection is achieved here thanks to the use of strain β3914 (Le Roux et al., 2007), a RP4+ and DAP-auxotroph strain, and thanks to a protocol recently optimized for other *Vibrio* species (Morot et al., 2021).

In order to study the ecophysiology of marine NCD, we next focused on the targeted deletion of genes. The *nifH* was the ideal candidate for gene deletion since it is found in only one copy in *V. diazotrophicus* NS1, it is conserved in diazotrophs and considered as a biomarker for diazotrophy and it codes the nitrogenase-reductase, and essential component of the nitrogenase multiprotein complex. Adapting an already existing protocol (Morot et al., 2021), we were able to perform a scarless deletion of the *nifH* gene in this strain. To our knowledge, this mutant represents the first marine NCD mutant described in literature, opening an avenue for a better characterization of their ecophysiology and role in the N and C biogeochemical cycles. Its nitrogen-fixation capability was subsequently tested thanks to a strain- and medium-adapted soft-gellan bioassay (Hashidoko et al., 2002; Hara et al., 2009). This soft-gellan assay allows the inoculated bacteria to “find” their best appropriate O_2_-tension for optimized growth, characterized by a visible ring formed within the semi-solid medium (Hashidoko et al., 2002). When nitrogen-free medium is used, the location of this growing ring corresponds to the delicate balance between O_2_ tension-necessary for aerobic respiration- and diazotrophy inhibition by O_2_. Additionally, gellan gum allows a complete transparency of the medium and better visualization of bacterial growth (Hashidoko et al., 2002), frequently allows faster growth and a higher bacterial diversity recovered (Tamaki et al., 2009; Delavat et al., 2012; Delavat et al., 2013) and, importantly, bacteria grown in soft-gellan show a much higher diazotrophic activity compared to the same bacteria grown in soft-agar (Hara et al., 2009).

When grown in a nitrogen-free soft-gellan medium, we clearly showed that, unlike *V. diazotrophicus* NS1 wild type, the Δ*nifH* mutant is unable to grow, while supplementing the medium with NO_3_^−^ or NH_4_^+^ as an organic nitrogen source allows its growth (Figure 3 and Figure S1). Moreover, *in trans* complementation of the *nifH* mutant restores its capacity to grow in a nitrogen-free medium, demonstrating that growth in this medium is dependent on an intact nitrogenase. We also show here that the developed soft-gellan bioassay is powerful to screen for diazotrophic activity and to phenotypically differentiate *V. diazotrophicus* mutants, *e.g*. those related to N_2_ fixation (catalytic or regulatory genes) or related to protection from O_2_. This experiment and the fact that no growth was obtained in the Δ*nifH* mutant even after prolonged incubation in this soft-gellan tubes also confirm the presence of only one copy of the *nifH* gene in the genome *V. diazotrophicus* NS1 and no paralog being able to partially complement this mutation (Figure 5B).

Because of the inhibition of diazotrophy by O_2_, nitrogen-fixation occurs at intermediate O_2_ tension. We demonstrated here with a nitrogen-free soft-gellan medium that *V. diazotrophicus* NS1 grows under microoxic conditions, ranging from 1 to 15 μM O_2_ to achieve a sufficient N_2_ fixation activity to sustain its growth (Figure 3). Decreasing the local O_2_ tension is therefore important, and can be achieved by different means (for a review, see (Gallon, 1992). As the ecophysiology of marine NCD is basically unknown, we tested whether *V. diazotrophicus* NS1 is able to produce a biofilm, which would restrict the diffusion of O_2_ in the cells. Indeed, this strain produces more biofilm when grown in a nitrogen-limited medium, compared with the biofilm formed in LB-rich medium (Figure 4A). This biofilm formation decreases when NO_3_^−^ or NH_4_^+^ is added as an alternative bioavailable nitrogen source. These results were similar for the Δ*nifH* mutant and the Δ*nifH* complemented strain (Figure 4A), demonstrating that biofilm production is not dependent on the presence of an intact nitrogenase and that both processes are genetically decoupled, at least at this genetic hierarchy. The thickness of the biofilm was quantified using flow cells, showing that *V. diazotrophicus* NS1 produces a substantial biofilm after 24 hours of growth (Figure 4BCD). The thickness of the biofilm measured in this experiment (on average of 26.65 μm) probably serves as a barrier for O_2_-diffusion. Indeed, this value is coherent with previous data, showing the rapid decrease in O_2_ in thick biofilms, which ultimately turn anoxic (Stewart et al., 2016). Thus, *V. diazotrophicus* NS1 produces biofilm when grown under nitrogen-limited condition, which might restrict O_2_ diffusion, thus help solving the dilemma of nitrogenase inhibition by O_2_. This behaviour is analog to the one of the terrestrial strain *Pseudomonas stutzeri* A1501, which produces biofilm under nitrogen-starved conditions (Wang et al., 2017). Marine non cyanobacterial diazotrophs are in contrast less studied. Cell aggregates of 1-4 mm in diameter were observed for the marine diazotroph *P. stutzeri* BAL361 when grown in nitrogen-free media under oxic condition, which was hypothesized to be an adaptive mechanism for *P. stutzeri* BAL361 to fix N_2_ despite high surrounding O_2_-tension (Bentzon-Tilia et al., 2015). Cell aggregation and exopolysaccharide production is also suspected to play a role in the diazotrophic activity of the marine NCD *Sagittula castanea*, even though cell aggregation was also observed under anoxic conditions in this strain (Martinez-Perez et al., 2018). In this study, we showed that *V. diazotrophicus* cells grown under microaerobic conditions in nitrogen-free media don’t form aggregates (Figure S2), and demonstrated an increase in biofilm density and biofilm thickness of a marine NCD when grown under nitrogen-limited conditions as compared to LB-grown culture, constituting an additional level of evidence that biofilm formation is important for these bacteria when grown under oxic condition. This hypothesis is reinforced by the recent demonstration of the high prevalence of marine NCD in high size fractions of marine metagenomic samples (Delmont et al., 2022), suggesting NCD aggregation on particles and planktonic aggregates (Farnelid et al., 2019; Geisler et al., 2019).

*V. diazotrophicus* is a bacterial species which has been isolated from different marine environments, including sea water (Guerinot et al., 1982), gastrointestinal tractus of sea urchin (Guerinot and Patriquin, 1981) or deep see sediments (Castillo et al., 2018), and from enriched-phytoplankton exometabolites (Fu et al., 2020). However, this species is barely detected from metagenomes coming from *Tara ocean* expeditions (Table S5), and is only distantly related to newly discovered MAGs (Delmont et al., 2022). This almost absent distribution from those metagenomic samples despite their effective presence in various marine environments suggests that additional NCD live in alternative ecological niches that have been overlooked by most metagenomic studies, like those from macroorganisms (*e.g*. see urchins) or those from deep see sediment (Castillo et al., 2018).

In conclusion, this study allowed unveiling the ecophysiology of a marine NCD belonging to the species *V. diazotrophicus*, which might serve as an ideal model to decipher their ecological importance related to the N and C biogeochemical cycles and their genetic and physiological adaptations to their environments, including the O_2_-tension and the presence of dissolved inorganic nitrogen.

## Acknowledgments

The authors would like to thank Corentin Ramaugé-Parra for his help in the initial optimization of the genetic tools. The authors acknowledge the MicroScope plateform (https://mage.genoscope.cns.fr/microscope/home/index.php) for providing access to their plateform. This work was supported by the “Connect Talent” EpiAlg project awarded to Leila Tirichine, and by the “Rising Star” SMIDIDI project awarded to François Delavat, both from the French Pays de la Loire Region.

---

*Supplementary Figure S1*

Soft-gellan assay demonstrating bacterial growth under appropriate microoxic conditions. The “+” and “−” indicate the presence or absence of NO_3_^−^, respectively. White arrows indicate the location of the growing ring. Note the absence of ring in the Δ*nifH* mutant and in the presence of NH_4_^+^.

*Supplementary Figure S2*

Microscopy picture of *V. diazotrophicus* NS1 pFD086 cells grown under microoxic condition in a nitrogen-free MDV soft-gellan medium. Cells from the growing ring were pipetted with a syringe and a needle, and observed by epifluorescence microscopy. Please note that cells are isolated and don’t form aggregates.

*Supplementary Figure S3*

(A) View of the top of the biofilm produced by *V. diazotrophicus* NS1 pFD086 after 48 hours of incubation in a flow cell. (B) three-dimensional representation showing the thickness of the biofilm produced after 48 hours.

