## Supplementary figures and images for "Genetic and physiological insights into the diazotrophic activity of a non-cyanobacterial marine diazotroph"

### Figure S1

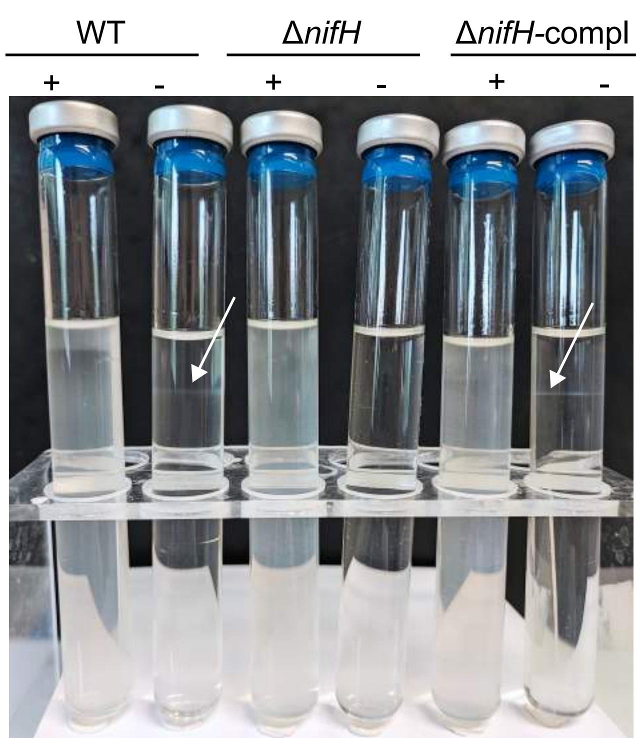

### Figure S2

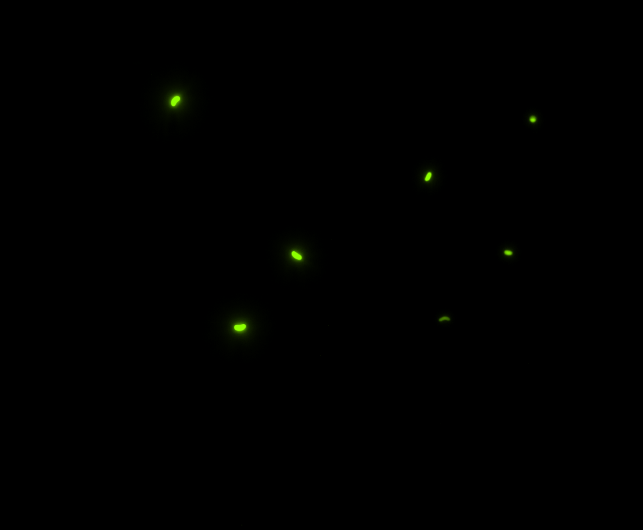

### Figure S3

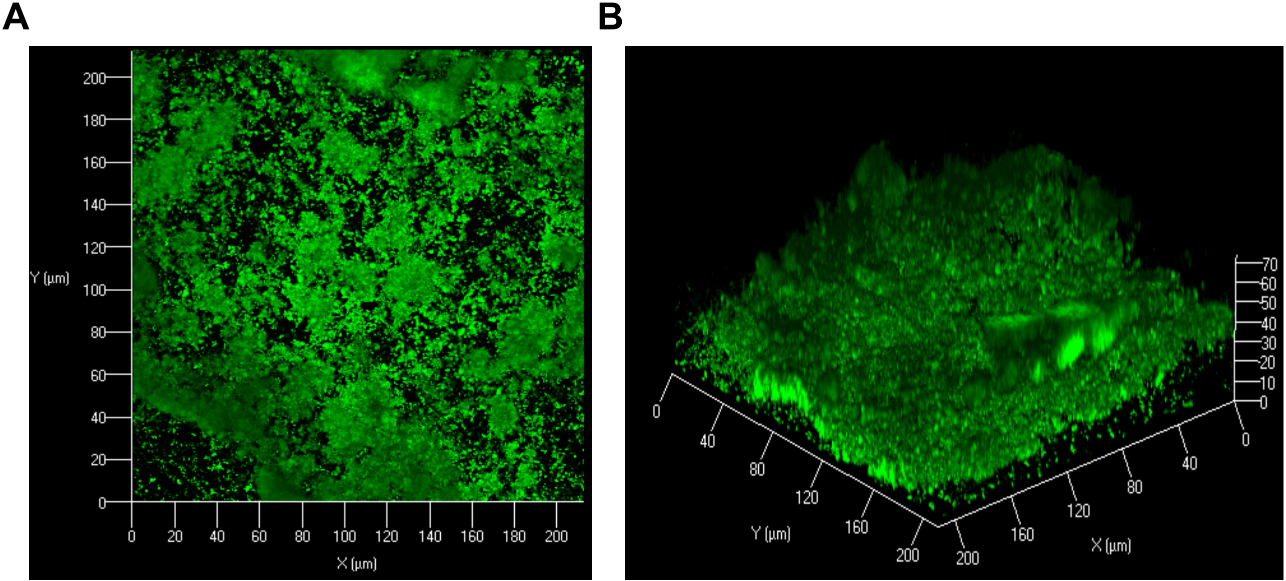
