## Supplementary material for "Genetic and physiological insights into the diazotrophic activity of a non-cyanobacterial marine diazotroph": Table S2

| Strain collection number (US2B lab collection) | Strain name | Characteristics | References |
| --- | --- | --- | --- |
| *Escherichia coli* | | | |
| 204 | DH5α λpir | *sup E44, ΔlacU169 (ΦlacZΔM15), recA1, endA1, hsdR17, thi-1, gyrA96, relA1, λpir phage lysogen* | Lab collection |
| 206 | GEB883 + pEVS104 | Strain GEB883 {Nguyen, 2018 #1059} containing pEVS104 {Stabb, 2002 #1060} | {Morot, 2021 #1321} |
| 207 | β-3914 | F^-^, RP4-2-Tc::Mu, Δ*dapA*::(*erm-pir*)  *gyrA462 zei-298::*Tn*10*, Km^R^, Em^R^, Tc^R^, DAP^-^ | {Le Roux, 2007 #1046} |
| *Vibrio diazotrophicus* | | | |
| 295 | *V. diazotrophicus* NBRC 103148 | Strain isolated from urchin gut | {Guerinot, 1981 #1280} |
| 301 | *V. diazotrophicus* pFD086 | Derivative of 295 containing pFD085 | This study |
| 357 | *V. diazotrophicus* Δ*nifH* | Derivative of strain 295 in which the *nifH* gene has been deleted | This study |
| 375 | *V. diazotrophicus* Δ*nifH* pFD120 | Derivative of strain 357 containing pFD120 | This study |
| 376 | *V. diazotrophicus* pFD085 | Derivative of strain 295 containing pFD085 | This study |
| 377 | *V. diazotrophicus* Δ*nifH* pFD085 | Derivative of strain 357 containing pFD085 | This study |

Table S1. Strains used in this study.
