## Supplementary material for "Genetic and physiological insights into the diazotrophic activity of a non-cyanobacterial marine diazotroph": Table S2

| Plasmid name | Plasmid characteristics | Reference |
| --- | --- | --- |
| pGEM-T | Cloning vector. *lacZ*. Amp^R^ | Promega |
| pLP12 | Suicide plasmid used for targeted deletion of chromosomal genes of *Vibrio* species. *oriT*_RP4_, *oriV*_R6K_, P*_BAD_*-*vmi480*. Cm^R^ | {Luo, 2015 #1224} |
| pEVS104 | Conjugative helper plasmid. oriV_R6K_ oriT_RP4_. Km^R^ | {Stabb, 2002 #1060} |
| pFD085 | Replicative plasmid for *Vibrio*, containing a promoterless *gfp* gene. Trim^R^ | {Morot, 2021 #1321} |
| pFD086 | Derivative of pFD085 containing the P*_lac_* promoter upstream of the *gfp* gene of pFD085 Trim^R^ | {Morot, 2021 #1321} |
| pFD114 | Derivative of pLP12 containing the upstream and downstream fragments of the *nifH* gene of *V. diazotrophicus*. Insertion performed using SmaI + EcoRI | This study |
| pFD120 | Derivative of pFD086 in which the P*_lac_*-*gfp* fragment has been replaced by the P*_nifH_*-*nifH* fragment of *V. diazotrophicus*, using XhoI + BamHI | This study |

Table S2. Plasmids used in this study. All *in silico* plasmid sequences and maps are available upon request
