## Supplementary material for "Genetic and physiological insights into the diazotrophic activity of a non-cyanobacterial marine diazotroph": Table S3

| Primer number | Primer name | Primer sequence | Target |
| --- | --- | --- | --- |
| 210204 | PnifH-R-PstI-XhoI | CTCGAGCTGCAGcgtttcttaattcgctagttc | Primer annealing 285 bp upstream of the ATG of *nifH* from *V. diazotrophicus* (strain 295). Used in combination with 210505 to construct plasmid pFD120 |
| 210210 | V.dia-up-nifH-F-SmaI | ATGCCCCGGGcgaggatgatgcgagaagac | Forward primer annealing 763 bp downstream of the stop codon of *nifH* from *V. diazotrophicus* (strain 295), facing towards *nifH*. Used in combination with 210211, 210212 and 210213 to construct plasmid pFD114 |
| 210211 | V.dia-up-nifH-R | Gcgtaagttaagcactgattc | Reverse primer annealing around the stop codon of nifH from *V. diazotrophicus* (strain 295), facing outwards. Used in combination with 210210, 210212 and 210213 to construct plasmid pFD114 |
| 210212 | V.dia-dn-nifH-F-rc-up | Gaatcagtgcttaacttacgccacattgacgaattgccatg | Forward primer annealing around the ATG of *nifH* from *V. diazotrophicus* (strain 295), facing outwards. Used in combination with 210210, 210211 and 210213 to construct plasmid pFD114 |
| 210213 | V. dia-dn-nifH-R-EcoRI | ATGCGAATTCgcagtcggttgaactgaaag | Reverse primer annealing 727 bp upstream of the ATG of *nifH* from *V. diazotrophicus* (strain 295), facing inwards. Used in combination with 210210, 210211 and 210212 to construct plasmid pFD114 |
| 210501 | vibrio-Dell-nifH-F | ccatcacctgaccattgag | Primer annealing 807 bp downstream of the *nifH* gene of *V. diazotrophicus*, facing inwards. Used in combination with 210502 to test for the deletion of *nifH* in this strain |
| 210502 | Vibrio-Dell-nifH-R | gtctgcaaagctatcttgaag | Primer annealing 785 bp upstream of the *nifH* gene of *V. diazotrophicus*, facing inwards. Used in combination with 210501 to test for the deletion of *nifH* in this strain |
| 210505 | stop-BamHI-nifH_vibrio-F | ATGCGGATCCccaaataagaagaatcagtgc | Primer annealing 32 bp downstream of the stop codon of *nifH* from *V. diazotrophicus* (strain 295). Used in combination with 210204 to construct plasmid pFD120 |

Table S3. Primers used in this study
