## Supplementary material for "Genetic and physiological insights into the diazotrophic activity of a non-cyanobacterial marine diazotroph": Table S4

**ANIb value computed with PyANI using "anvi-compute-genome-similarity" from Anvi'o**

| | HBD_Gamma_02 | *V. diazotrophicus* NS1 |
|---|---|---|
| HBD_Gamma_02 | 1.0 | 0.687493790322581 |
| *V. diazotrophicus* NS1 | 0.6861198461538462 | 1.0 |

| | *V. diazotrophicus* NS1 | *V. diazotrophicus* 60.18M | *V. diazotrophicus* 60.27F | *V. diazotrophicus* 60.6B |
|---|---|---|---|---|
| *V. diazotrophicus* NS1 | 1.0 | 0.9752714603256655 | 0.976739213229767 | 0.9421276906552095 |
| *V. diazotrophicus* 60.18M | 0.9747959229576008 | 1.0 | 0.9765777350533367 | 0.9400872675521821 |
| *V. diazotrophicus* 60.27F | 0.97671442182001 | 0.9763992024844721 | 1.0 | 0.9416596740858506 |
| *V. diazotrophicus* 60.6B | 0.9418917185821698 | 0.9401380649526389 | 0.9415518022793533 | 1.0 |
| *V. diazotrophicus* 60.6F | 0.9743143829355002 | 0.9752033937562941 | 0.9767368589108911 | 0.9415780806749275 |
| *V. diazotrophicus* 65.10M | 0.9738431454081632 | 0.9733662619888945 | 0.9754824803536346 | 0.9393013855259617 |
| *V. diazotrophicus* 65.7M | 0.9733284013174565 | 0.9730305924781041 | 0.9744740159441954 | 0.9397564740502942 |
| *V. diazotrophicus* 99A | 0.9651257999489665 | 0.9646872820250576 | 0.9672845241654211 | 0.9517851016684047 |
| *V. diazotrophicus* HF9B | 0.9029173517465475 | 0.9041391805974239 | 0.9033061027027028 | 0.8985644718909711 |

| *V. diazotrophicus* 60.6F | *V. diazotrophicus* 65.10M | *V. diazotrophicus* 65.7M | *V. diazotrophicus* 99A | *V. diazotrophicus* HF9B |
|---|---|---|---|---|
| 0.9747018343949044 | 0.9740273962070732 | 0.9731189353612169 | 0.9662374389930645 | 0.9031162761506276 |
| 0.9751769399044506 | 0.9735739766970618 | 0.972487849434738 | 0.9652297167138811 | 0.9038819293325855 |
| 0.9766205573445628 | 0.9757248350566222 | 0.9746162873505977 | 0.9680632016008002 | 0.9035468236277515 |
| 0.9418244341862306 | 0.9402812242686891 | 0.9399978016085793 | 0.9529524461942256 | 0.8983564387963775 |
| 1.0 | 0.973914864522664 | 0.9731384256861728 | 0.9663636737331956 | 0.9031073251847641 |
| 0.9735753129732458 | 1.0 | 0.9744342049643647 | 0.9741496490372897 | 0.9028888171444477 |
| 0.9733997692507128 | 0.9748823545432038 | 1.0 | 0.9675132430398797 | 0.903290241264559 |
| 0.9659991615067078 | 0.9738949744587692 | 0.9664535275080905 | 1.0 | 0.9015833099579241 |
| 0.9031158662613982 | 0.9034341267682264 | 0.9036073782467533 | 0.9022546803278688 | 1.0 |
