## Supplementary material for "Genetic and physiological insights into the diazotrophic activity of a non-cyanobacterial marine diazotroph": Table S5

**Vibrio diazotrophicus NS1 genome detection in the Tara ocena metagenomes.**

| PANGAEA sample id | Detection |
|---|---|
| TARA_B110001452 | 0,00318271 |
| TARA_B110001454 | 0,00403164 |
| TARA_B110001450 | 0,00248321 |
| TARA_B110000196 | 0,00506287 |
| TARA_B110001469 | 0,00316088 |
| TARA_B110000046 | 0,00489944 |
| TARA_B110000037 | 0,00232551 |
| TARA_B110000027 | 0,00165505 |
| TARA_B110000091,TARA_E | 0,00505587 |
| TARA_B110000090 | 0,00319479 |
| TARA_B110000116 | 0,00289252 |
| TARA_B110000114 | 0,00254277 |
| TARA_B110000208 | 0,00276259 |
| TARA_B110000211 | 0,00606887 |
| TARA_B110000503 | 0,00326516 |
| TARA_B110000240 | 0,00375312 |
| TARA_B110000238 | 0,00205821 |
| TARA_B110000261 | 0,00143736 |
| TARA_B110000263 | 0,00197788 |
| TARA_B110000259 | 0,00165971 |
| TARA_B110000285 | 0,00217374 |
| TARA_B110000977 | 0,00193294 |
| TARA_B110000967 | 0,00233759 |
| TARA_B110000971 | 0,00196495 |
| TARA_B110000305 | 0,00183734 |
| TARA_B110000908 | 0,00290757 |
| TARA_B110000914 | 0,00267928 |
| TARA_B110000902 | 0,00305595 |
| TARA_B110000881 | 0,00290863 |
| TARA_B110000879 | 0,00231576 |
| TARA_B110000495 | 0,00239228 |
| TARA_B110000483 | 0,00150816 |
| TARA_B110000858 | 0,00135999 |
| TARA_B110000467 | 0,0035208 |
| TARA_B110000459 | 0,00186236 |
| TARA_B110000438 | 0,00316299 |
| TARA_B110000444 | 0,00209849 |
| TARA_B100000035 | 0,00608795 |
| TARA_Y100000015,TARA_Y | 0,00434006 |
| TARA_Y100000310,TARA_E | 0,03568279 |
| TARA_B100000073 | 0,00534224 |
| TARA_Y100000296,TARA_Y | 0,01265982 |
| TARA_Y100000287,TARA_Y | 0,01640085 |
| TARA_B100000085 | 0,00358905 |
| TARA_Y100000034,TARA_Y | 0,01501882 |

| | |
|---|---|
| TARA_Y100000033 | 0,00365815 |
| TARA_B100000287 | 0,00552941 |
| TARA_B100000282,TARA_Y | 0,0068521 |
| TARA_B100000131 | 0,00332451 |
| TARA_B100000123 | 0,00296226 |
| TARA_B100000161 | 0,0042612 |
| TARA_B100000242,TARA_Y | 0,00531172 |
| TARA_B100000214 | 0,00304493 |
| TARA_B100000212 | 0,00486616 |
| TARA_B100000378 | 0,00344576 |
| TARA_B000000609 | 0,0022776 |
| TARA_B000000565 | 0,00653796 |
| TARA_B000000557 | 0,00398988 |
| TARA_B000000532 | 0,00340506 |
| TARA_Y100000401,TARA_E | 0,00526742 |
| TARA_B100000408 | 0,00535221 |
| TARA_B100000401 | 0,0045516 |
| TARA_Y100000361,TARA_E | 0,00509615 |
| TARA_B000000460 | 0,0060888 |
| TARA_Y100000356,TARA_E | 0,00319861 |
| TARA_S200000501 | 0,00526615 |
| TARA_A100000164 | 0,00525958 |
| TARA_E500000081 | 0,00814786 |
| TARA_E500000075 | 0,00445452 |
| TARA_E500000331 | 0,01026839 |
| TARA_E500000178 | 0,00752891 |
| TARA_A100001011 | 0,0040982 |
| TARA_A100001015 | 0,0075022 |
| TARA_A200000159 | 0,01314862 |
| TARA_A200000113 | 0,0131323 |
| TARA_X000001036 | 0,00525216 |
| TARA_X000000950 | 0,01131488 |
| TARA_B100001939 | 0,00912143 |
| TARA_B100002052 | 0,00396678 |
| TARA_B100002049 | 0,00950933 |
| TARA_B100002051 | 0,00385422 |
| TARA_B100001146 | 0,00498592 |
| TARA_B100001142 | 0,00183713 |
| TARA_B100001167 | 0,00350066 |
| TARA_B100001540 | 0,00216653 |
| TARA_B100001750 | 0,00571319 |
| TARA_B100001741 | 0,00290418 |
| TARA_B100001765 | 0,00400069 |
| TARA_B100001758 | 0,00171843 |
| TARA_B100001778 | 0,00411495 |
| TARA_B100001769 | 0,00311488 |
| TARA_B100001559 | 0,00375481 |
| TARA_B100001564 | 0,0047818 |

| | |
|---|---|
| TARA_B100001179 | 0,00462325 |
| TARA_B100001175 | 0,00283995 |
| TARA_B100001173 | 0,00224157 |
| TARA_B110000008 | 0,02365739 |
| TARA_B110000014 | 0,02143638 |
| TARA_B110000003 | 0,01623616 |
| TARA_X000000368 | 0,00380865 |
| TARA_Y200000002 | 0,00451154 |
| TARA_B100000927 | 0,00600443 |
| TARA_B100000929 | 0,01398865 |
| TARA_B100000925 | 0,0068451 |
| TARA_B100001250 | 0,00360813 |
| TARA_B100001245 | 0,00657972 |
| TARA_B100001248 | 0,00301546 |
| TARA_B100001094 | 0,00366154 |
| TARA_B100001105 | 0,01752704 |
| TARA_B100001093 | 0,00554722 |
| TARA_B100001964 | 0,00457025 |
| TARA_B100001971 | 0,00704223 |
| TARA_B100001287 | 0,00404648 |
| TARA_B100001996 | 0,00517691 |
| TARA_B100002003 | 0,01279612 |
| TARA_B100001989 | 0,00360453 |
| TARA_B100002019 | 0,00700302 |
| TARA_A100001388 | 0,00300699 |
| TARA_A100001037 | 0,01011323 |
| TARA_A100001035 | 0,00836682 |
| TARA_A100001234 | 0,00365815 |
| TARA_B100000029 | 0,00632196 |
| TARA_Y100000004,B10000 | 0,00395555 |
| TARA_B000000477 | 0,00233102 |
| TARA_B000000475 | 0,00180152 |
| TARA_Y100000389,TARA_E | 0,00408654 |
| TARA_Y100000746,TARA_E | 0,0083895 |
| TARA_Y100000758,TARA_E | 0,01006533 |
| TARA_B100000475,TARA_Y | 0,00780214 |
| TARA_B100000446,TARA_Y | 0,01287836 |
| TARA_B100000459,TARA_Y | 0,01052636 |
| TARA_B100000427 | 0,0043195 |
| TARA_B100000508 | 0,01669676 |
| TARA_B100000424 | 0,00426862 |
| TARA_B100000519,TARA_Y | 0,00776568 |
| TARA_B100000749,TARA_Y | 0,02102855 |
| TARA_Y100000813,TARA_E | 0,00547557 |
| TARA_Y100000996,TARA_E | 0,00530049 |
| TARA_Y100001001,TARA_E | 0,01099926 |
| TARA_B100000524,TARA_Y | 0,00643409 |
| TARA_B100000767 | 0,00961913 |

| | |
|---|---|
| TARA_B100000768 | 0,00621195 |
| TARA_B100000780 | 0,00236112 |
| TARA_B100000795 | 0,00250123 |
| TARA_B100000809 | 0,00433921 |
| TARA_B100000787 | 0,00188567 |
| TARA_B100000959,TARA_E | 0,00851329 |
| TARA_B100000963 | 0,01166654 |
| TARA_B100000902 | 0,00833015 |
| TARA_B100000953 | 0,01229418 |
| TARA_B100000900 | 0,00616659 |
| TARA_B100001113 | 0,00793229 |
| TARA_B100001079 | 0,00748355 |
| TARA_B100001109 | 0,00712087 |
| TARA_B100000579 | 0,00562586 |
| TARA_B100000586 | 0,0065157 |
| TARA_B100000575 | 0,00488503 |
| TARA_B100000945 | 0,00383409 |
| TARA_B100000949 | 0,00463512 |
| TARA_B100000941 | 0,00241114 |
| TARA_Y100001968,TARA_E | 0,01306002 |
| TARA_Y100001951,TARA_E | 0,01900065 |
| TARA_Y100001978,TARA_Y | 0,00815507 |
| TARA_Y100001956,TARA_Y | 0,08587249 |
| TARA_Y100001958,TARA_E | 0,01795352 |
| TARA_Y100001938,TARA_Y | 0,03368371 |
| TARA_Y100001933,TARA_E | 0,01337055 |
| TARA_Y100000589,TARA_E | 0,02935086 |
| TARA_Y100000592,TARA_Y | 0,07681931 |
| TARA_B100000614 | 0,00631391 |
| TARA_B100000609 | 0,00421542 |
| TARA_B100001059 | 0,06052932 |
| TARA_B100001063 | 0,08928921 |
| TARA_B100001057 | 0,03355971 |
| TARA_B100000989 | 0,00264982 |
| TARA_B100001029 | 0,00200098 |
| TARA_B100001013 | 0,00506435 |
| TARA_B100001027 | 0,00180449 |
| TARA_B100000886 | 0,00400324 |

| PANGAEA sample id | Detection |
|---|---|
| TARA_A100000393 | 0,00392608 |
| TARA_A100000394 | 0,00208959 |
| TARA_A100000400 | 0,00207136 |
| TARA_A100000402 | 0,00211439 |
| TARA_A100000534 | 0,0039388 |
| TARA_A100000536 | 0,00115714 |
| TARA_A100000538 | 0,00433603 |
| TARA_A100000539 | 0,00258835 |
| TARA_A100000549 | 0,00837255 |
| TARA_A100000550 | 0,00442569 |
| TARA_A100000552 | 0,00506796 |
| TARA_A100000595 | 0,00264155 |
| TARA_A100000596 | 0,00267398 |
| TARA_A100000598 | 0,00334317 |
| TARA_A100000759 | 0,00608223 |
| TARA_A100001641 | 0,00802047 |
| TARA_A100001647 | 0,00488248 |
| TARA_A100001649 | 0,00855293 |
| TARA_A200000123 | 0,0201828 |
| TARA_A200000139 | 0,04311928 |
| TARA_E400007200 | 0,00332812 |
| TARA_N000000006 | 0,00527187 |
| TARA_N000000011 | 0,01048693 |
| TARA_N000000029 | 0,00470782 |
| TARA_N000000063 | 0,01023045 |
| TARA_N000000071 | 0,00613607 |
| TARA_N000000076 | 0,01046955 |
| TARA_N000000077 | 0,03062755 |
| TARA_N000000113 | 0,00674166 |
| TARA_N000000184 | 0,01499572 |
| TARA_N000000189 | 0,00680228 |
| TARA_N000000196 | 0,00318144 |
| TARA_N000000202 | 0,03397284 |
| TARA_N000000203 | 0,01417561 |
| TARA_N000000214 | 0,0072453 |
| TARA_N000000267 | 0,00736167 |
| TARA_N000000268 | 0,00708632 |
| TARA_N000000269 | 0,01099227 |
| TARA_N000000270 | 0,02933624 |
| TARA_N000000278 | 0,00577296 |
| TARA_N000000284 | 0,01889551 |
| TARA_N000000311 | 0,00599744 |
| TARA_N000000313 | 0,02518526 |
| TARA_N000000316 | 0,00435998 |
| TARA_N000000522 | 0,00339277 |
| TARA_N000000526 | 0,00269179 |
| TARA_N000000539 | 0,00729871 |

| | |
|---|---|
| TARA_N000000581 | 0,00265448 |
| TARA_N000000589 | 0,00372153 |
| TARA_N000000598 | 0,00433433 |
| TARA_N000000616 | 0,00359456 |
| TARA_N000000625 | 0,002908 |
| TARA_N000000662 | 0,00249657 |
| TARA_N000000666 | 0,00804527 |
| TARA_N000000674 | 0,00356531 |
| TARA_N000000678 | 0,00396106 |
| TARA_N000000682 | 0,00396508 |
| TARA_N000000705 | 0,00088348 |
| TARA_N000000709 | 0,00163703 |
| TARA_N000000722 | 0,00270366 |
| TARA_N000000742 | 0,00119529 |
| TARA_N000000746 | 0,00170359 |
| TARA_N000000750 | 0,00329717 |
| TARA_N000000756 | 0,00124701 |
| TARA_N000000759 | 0,00133752 |
| TARA_N000000790 | 0,01027242 |
| TARA_N000000793 | 0,00145135 |
| TARA_N000000805 | 0,0021023 |
| TARA_N000000839 | 0,0022651 |
| TARA_N000000843 | 0,00128262 |
| TARA_N000000852 | 0,00382497 |
| TARA_N000000863 | 0,00301695 |
| TARA_N000000871 | 0,00408379 |
| TARA_N000000879 | 0,00687902 |
| TARA_N000000928 | 0,00305871 |
| TARA_N000000968 | 0,00269582 |
| TARA_N000000981 | 0,00564557 |
| TARA_N000001006 | 0,00139391 |
| TARA_N000001017 | 0,00104161 |
| TARA_N000001024 | 0,0019287 |
| TARA_N000001028 | 0,00233102 |
| TARA_N000001288 | 0,00240118 |
| TARA_N000001290 | 0,01028302 |
| TARA_N000001292 | 0,00709332 |
| TARA_N000001296 | 0,00864662 |
| TARA_N000001299 | 0,00731991 |
| TARA_N000001301 | 0,0021057 |
| TARA_N000001303 | 0,00160587 |
| TARA_N000001305 | 0,00923059 |
| TARA_N000001306 | 0,00472096 |
| TARA_N000001309 | 0,00397059 |
| TARA_N000001362 | 0,00073193 |
| TARA_N000001372 | 0,0045251 |
| TARA_N000001374 | 0,00692353 |
| TARA_N000001380 | 0,00190093 |

| | |
|---|---|
| TARA_N000001382 | 0,00164594 |
| TARA_N000001386 | 0,00439411 |
| TARA_N000001394 | 0,00126058 |
| TARA_N000001398 | 0,00199187 |
| TARA_N000001428 | 0,01103127 |
| TARA_N000001436 | 0,00714546 |
| TARA_N000001438 | 0,00184285 |
| TARA_N000001442 | 0,00078852 |
| TARA_N000001478 | 0,00891773 |
| TARA_N000001491 | 0,00773325 |
| TARA_N000001499 | 0,04640839 |
| TARA_N000001503 | 0,0289159 |
| TARA_N000001510 | 0,00290418 |
| TARA_N000001514 | 0,00528035 |
| TARA_N000001578 | 0,00598451 |
| TARA_N000001582 | 0,00183777 |
| TARA_N000001586 | 0,00310152 |
| TARA_N000001604 | 0,0027753 |
| TARA_N000001608 | 0,00216823 |
| TARA_N000001612 | 0,00123069 |
| TARA_N000001616 | 0,00431695 |
| TARA_N000001620 | 0,01450776 |
| TARA_N000001646 | 0,00309792 |
| TARA_N000001650 | 0,00297922 |
| TARA_N000001654 | 0,00397695 |
| TARA_N000001658 | 0,00451514 |
| TARA_N000001662 | 0,00738329 |
| TARA_N000001727 | 0,00377092 |
| TARA_N000001730 | 0,00346357 |
| TARA_N000001734 | 0,00456072 |
| TARA_N000001738 | 0,00410859 |
| TARA_N000001742 | 0,01440856 |
| TARA_N000001746 | 0,0033508 |
| TARA_N000001750 | 0,00485832 |
| TARA_N000001754 | 0,00517309 |
| TARA_N000001758 | 0,0027628 |
| TARA_N000001762 | 0,00479876 |
| TARA_N000001810 | 0,00488439 |
| TARA_N000001812 | 0,00222503 |
| TARA_N000001823 | 0,00259661 |
| TARA_N000001826 | 0,02134672 |
| TARA_N000001937 | 0,00509912 |
| TARA_N000001938 | 0,00235094 |
| TARA_N000001941 | 0,00373361 |
| TARA_N000001942 | 0,00929546 |
| TARA_N000001943 | 0,05085062 |
| TARA_N000001972 | 0,00572527 |
| TARA_N000001992 | 0,00515317 |

| | |
|---|---|
| TARA_N000001994 | 0,00495625 |
| TARA_N000001996 | 0,01105013 |
| TARA_N000001998 | 0,0196533 |
| TARA_N000002017 | 0,00666111 |
| TARA_N000002019 | 0,00348985 |
| TARA_N000002021 | 0,00506393 |
| TARA_N000002024 | 0,00685358 |
| TARA_N000002025 | 0,01541774 |
| TARA_N000002036 | 0,00434959 |
| TARA_N000002037 | 0,00542851 |
| TARA_N000002039 | 0,01467373 |
| TARA_N000002041 | 0,00245587 |
| TARA_N000002043 | 0,00530451 |
| TARA_N000002101 | 0,00153804 |
| TARA_N000002103 | 0,00186617 |
| TARA_N000002106 | 0,00183056 |
| TARA_N000002107 | 0,00145474 |
| TARA_N000002109 | 0,00511968 |
| TARA_N000002111 | 0,00269603 |
| TARA_N000002115 | 0,00261188 |
| TARA_N000002117 | 0,00349557 |
| TARA_N000002119 | 0,0016648 |
| TARA_N000002137 | 0,00243658 |
| TARA_N000002139 | 0,00430551 |
| TARA_N000002141 | 0,00293725 |
| TARA_N000002175 | 0,00476866 |
| TARA_N000002179 | 0,0017144 |
| TARA_N000002185 | 0,00267716 |
| TARA_N000002193 | 0,00401066 |
| TARA_N000002285 | 0,00422157 |
| TARA_N000002289 | 0,00988685 |
| TARA_N000002293 | 0,04598764 |
| TARA_N000002297 | 0,00216568 |
| TARA_N000002301 | 0,01121038 |
| TARA_N000002348 | 0,00339574 |
| TARA_N000002352 | 0,00428367 |
| TARA_N000002356 | 0,00228523 |
| TARA_N000002360 | 0,00364247 |
| TARA_N000002364 | 0,00304069 |
| TARA_N000002400 | 0,01276602 |
| TARA_N000002404 | 0,00980736 |
| TARA_N000002412 | 0,00332685 |
| TARA_N000002416 | 0,00345106 |
| TARA_N000002420 | 0,00387521 |
| TARA_N000002464 | 0,00353585 |
| TARA_N000002695 | 0,00219769 |
| TARA_N000002697 | 0,00201158 |
| TARA_N000002699 | 0,00199971 |

| | |
|---|---|
| TARA_N000002701 | 0,00123281 |
| TARA_N000002703 | 0,00239079 |
| TARA_N000002737 | 0,00310004 |
| TARA_N000002741 | 0,00256567 |
| TARA_N000002745 | 0,00218264 |
| TARA_N000002749 | 0,00293534 |
| TARA_N000002753 | 0,00786001 |
| TARA_N000002773 | 0,00184625 |
| TARA_N000002777 | 0,0033614 |
| TARA_N000002781 | 0,0014274 |
| TARA_N000002785 | 0,0028478 |
| TARA_N000002789 | 0,00110499 |
| TARA_N000002923 | 0,00424446 |
| TARA_N000002925 | 0,00412152 |
| TARA_N000002927 | 0,01105183 |
| TARA_N000002929 | 0,0096717 |
| TARA_N000002931 | 0,00720312 |
| TARA_N000002959 | 0,00478434 |
| TARA_N000002961 | 0,00412024 |
| TARA_N000002963 | 0,00282596 |
| TARA_N000002965 | 0,00211629 |
| TARA_N000002967 | 0,00393922 |
| TARA_N000003005 | 0,003666 |
| TARA_N000003007 | 0,01022918 |
| TARA_N000003035 | 0,00455732 |
| TARA_N000003037 | 0,00600719 |
| TARA_N000003039 | 0,01103763 |
| TARA_N000003044 | 0,00525406 |
| TARA_N000003081 | 0,00395491 |
| TARA_N000003083 | 0,00327894 |
| TARA_N000003087 | 0,00218137 |
| TARA_N000003089 | 0,01038158 |
| TARA_N000003141 | 0,00341185 |
| TARA_N000003153 | 0,00291414 |
| TARA_N000003175 | 0,00508025 |
| TARA_N000003187 | 0,00262565 |
| TARA_N000003191 | 0,00217374 |
| TARA_N000003215 | 0,00287408 |
| TARA_N000003219 | 0,01579187 |
| TARA_N000003223 | 0,00238549 |
| TARA_N000003227 | 0,00276386 |
| TARA_N000003231 | 0,04245836 |
| TARA_N000003240 | 0,00585669 |
| TARA_N000003246 | 0,00207602 |
| TARA_N000003253 | 0,00138945 |
| TARA_N000003257 | 0,00246774 |
| TARA_S400007202 | 0,00182335 |
| TARA_X000000323 | 0,00615218 |

| | |
|---|---|
| TARA_X000000325 | 0,00600337 |
| TARA_X000000357 | 0,00710391 |
| TARA_X000000954 | 0,00957504 |
| TARA_X000001006 | 0,042491 |
| TARA_X000001288 | 0,01063043 |
| TARA_X000001334 | 0,02478357 |
| TARA_X000001338 | 0,01296781 |
